# Rapid Evolution of Parasite Resistance via Improved Recognition and Accelerated Immune Activation and Deactivation

**DOI:** 10.1101/2020.07.03.186569

**Authors:** Amanda K. Hund, Lauren E. Fuess, Mariah L. Kenney, Meghan F. Maciejewski, Joseph M. Marini, Kum Chuan Shim, Dan I. Bolnick

## Abstract

Closely related populations often differ in resistance to a given parasite, as measured by infection success or failure. Yet, the immunological mechanisms of these evolved differences are rarely specified. Does resistance evolve via changes to the host’s ability to recognize that an infection exists, actuate an effective immune response, or attenuate that response? We tested whether each of these phases of the host response contributed to threespine sticklebacks’ recently evolved resistance to their tapeworm *Schistocephalus solidus.* While marine stickleback and some susceptible lake fish permit fast-growing tapeworms, other lake populations are resistant and suppress tapeworm growth via a fibrosis response. We subjected lab-raised fish from three populations (susceptible marine ‘ancestors’, a susceptible lake, a resistant lake), to a novel immune challenge (injection of: 1) a saline control, 2) alum, a generalized pro-inflammatory adjuvant that causes fibrosis, 3) a tapeworm protein extract, and 4) a combination of alum and tapeworm protein). All three populations were capable of a robust fibrosis response to the alum treatments (but not the saline control). Yet, only the resistant population exhibited a fibrosis response to the tapeworm protein alone. Thus, these populations differed in their ability to recognize the tapeworm but shared an intact fibrosis pathway. However, the resistant population also initiated fibrosis faster, and was able to attenuate fibrosis, unlike the susceptible populations slow but longer-lasting response to alum. As fibrosis has presumed pathological side-effects, this difference may reflect adaptions to mitigate costs of immunity in the resistant population. Broadly, our results confirm that parasite detection, activation speed, and immune attenuation simultaneously contribute to adaptations to parasite infection in natural populations.

**IMPACT SUMMARY:** Dramatic variation in parasite resistance is common in nature, even to the same parasite, yet we are still working to understand the mechanisms of how such differences evolve. Many evolution studies focus on the broad outcomes of infection (infected or not) when studying this question, without specifying what part of the immune response has evolved. Here, we experimentally partition different sequential stages in the host immune response (recognition, actuation, attenuation), to evaluate which stage(s) underly the evolution of host resistance to infection. This study compares three populations of threespine stickleback that naturally differ in their exposure and their ability to resist infections of a freshwater tapeworm. These include a “resistant” lake population, a “susceptible” lake population, and an ancestral marine population that is rarely exposed to the tapeworm in nature, but is susceptible when exposed in the lab. The resistant population exhibits a fibrosis immune response to infection, which has previously been linked to suppressed tapeworm growth and viability. We injected different immune challenges directly into the site of infection (peritoneal cavity) and measured the subsequent fibrosis response through time. We found that all populations were capable of producing fibrosis in response to a general immune stimulant (alum). But, only the resistant population was able to recognize and respond to tapeworm protein alone. This population also responded faster than the others, within 24 hours, and attenuated its fibrosis by 90 days post-injections whereas the other populations exhibited a slower response that did not attenuate in the study time-frame. We concluded that variation in parasite recognition, an early phase in the host response, shapes the evolution of the initiation and resolution of the physical response to infection. Broadly, our results support that parasite detection mechanisms could play a key role in the rapid evolution of parasite resistance.

## INTRODUCTION

Parasites can be a major source of selection on their hosts. In nature, they are common and costly, reducing host life span and fecundity. Hosts have therefore evolved elaborate and varied defense strategies to combat parasites, and host-parasite relationships are often characterized by a co-evolutionary arms race with adaptations and counter-adaptations cycling or escalating through time (Carius *et al.* 2001; Schulte *et al.* 2010). Evidence suggests that this arms race can shift rapidly across space and time (Hoeksema & Forde 2008; Karvonen & Seehausen 2012; Fernandes *et al.* 2019), generating stochastic or deterministic variation between isolated populations (Papkou *et al.* 2016). Indeed, many closely-related host populations vary in their ability to resist infection, even to the same parasite (Roy & Kirchner 2000; Boots *et al.* 2009; Vale *et al.* 2011). While this is a common pattern, our understanding of the underlying mechanisms that drive such rapid evolution of parasite resistance is still limited, particularly in wild systems.

Most evolutionary studies focus on the broad outcomes of infection, such as infection rates, resistant/susceptible phenotypes, host mortality, or measures of parasite load. However, infection outcomes depend on a series of sequential and stepwise interactions between hosts and parasites (Hall *et al.* 2017). Understanding where in this step-wise chain of events variation is occurring is essential to understanding what selective pressures are shaping host-parasite evolution. For example, environmental, ecological, or behavioral factors can influence the degree to which hosts are exposed to parasites (Stutz *et al.* 2014; Barron *et al.* 2015). Once infected, host resistance depends on the host’s ability to (1) detect a parasite, (2) actuate the appropriate response at a level that will eliminate or control the parasite, then (3) attenuate that response (Hall *et al.* 2017). Each of these steps is often costly and can cause self-damage, such as the energetic cost of initiating and maintaining an activated state (Ganeshan & Chawla 2014), the risk of auto-immune responses (recognition), or oxidative stress from effector response itself (Viney *et al.* 2005). Therefore, hosts should avoid initiating immune responses unnecessarily, and when activated, must able to turn off or modulate this response at the appropriate time, not prematurely, but also not too late as to impose excess cost once the danger has passed (Khan *et al.* 2017; Armour *et al.* 2020). Variation at any stage of the response can lead to differences in infection outcomes and immunopathology side-effects, yet when we only examine the end result, we do not know at which step variation occurred. While breaking down and isolating different steps of the infection cycle can be challenging, it is crucial for understanding the evolution of parasite resistance. Each step influences selection on subsequent steps and likely entails very different genes, selective pressures, costs, and opportunities for parasite counter-adaptation.

Taking a stepwise approach, and experimentally breaking apart different phases of host-parasite interactions, has the potential to be a powerful strategy for understanding the causes of variation in infection outcomes, and for partitioning the genetic and environmental contributions to parasite resistance or susceptibility. To date, this approach has largely been confined to functional cell and molecular biology studies, but has the potential to greatly enhance our understanding of host-parasite coevolution at larger scales (Hall *et al.* 2017, 2019). By using this approach, and experimentally partitioning different phases of the host response to infection through time, we provide a detailed look at the evolution of resistance in closely related populations of threespine stickleback that vary in their response to the tapeworm parasite, *Schistocephalus solidus.*

The threespine stickleback, a small northern temperate fish, has become a model system for studying the process of local adaptation because of their repeated independent colonization of freshwater lakes from the ancestral marine population. In British Columbia, Canada, these colonization events occurred approximately 11,000 years ago, after deglaciation (Bell & Foster 1994). Within these freshwater environments, stickleback are an intermediate host to a tapeworm parasite, which rarely infects anadromous stickleback (Simmonds & Barber 2016). Infections of *S. solidus* can be quite costly for fish: decreasing antipredator responses (Giles 1983; Milinski 1985), swimming ability (Blake *et al.* 2006), lowering body condition and energy reserves (Tierney *et al.* 1996; Barber & Svensson 2003), and decreasing investment in reproduction (Schultz *et al.* 2006), though these effects can vary by population (MacNab *et al.* 2009). While there is evidence that *S. solidus* can suppress host immunity (Scharsack *et al.* 2004), some populations of freshwater stickleback are able to avoid that suppression and mount an effective resistance response to the parasite (Lohman *et al.* 2017; Weber *et al.* 2017b, *in prep*, Fuess *et al. in prep*). Certain populations can suppress tapeworm growth by an immune response involving extensive fibrotic tissue throughout the peritoneal cavity where infections occur. Fibrosis can even kill small tapeworms through the formation of granulomas that fully encase small parasites and have been observed to contain moribund or dead tapeworms (Weber *et al in prep*, *pers obs*). Surveys in natural populations show that infection rates and tapeworm size are negatively correlated with the presence of fibrosis (Weber *et al. in prep*). Likewise, fibrosis reduces cestode growth in experimental infections in the laboratory (Weber *et al. in prep*). Fibrosis rarely occurs spontaneously in laboratory stickleback without an immune challenge (and even then, only weakly, *pers obs*). Although fibrosis thus appears to be an evolutionary adaptive response to infections, it also represents costly pathology. In the field, both females and males are less likely to reproduce when they have fibrosis (controlling for their infection status) (De Lisle & Bolnick *in prep*). Given this mix of costs and benefits, evolution should act to minimize unnecessary initiation of fibrosis, while also maximizing the rapid onset of fibrosis against infection, as well as the rate of clearance after the infection has passed.

To understand how selection on and variation in different aspects of the fibrosis response may be generating among-population differences in parasite resistance, we need to tease apart where in the infection cycle variation is arising. To do this, we focused on three populations: 1) a “resistant” lake population where preliminary data had suggested low infection rates, small tapeworms, and evidence of common and severe fibrosis in nature, 2) a “susceptible” lake population with high infection rates, large tapeworms, and rare fibrosis in nature, and 3) an “ancestral” marine population with negligible exposure to the tapeworm, no fibrosis observed in nature, and high susceptibility to laboratory infections (Weber *et al.* 2017a). We first performed a field survey in both lake populations to confirm population-level differences in infection rates and the presence of fibrosis. We then performed a lab experiment that isolated different stages of the host response using four different injected immune challenges delivered directly into the site of *S. Solidus* infection. Our experimental design allows us to test several hypotheses for why parasite resistance varied among these populations: H1) variation in resistance is driven by ecological factors (i.e. lake differences) and all populations will mount similar responses to immune challenges in a common garden lab setting, H2) populations differ in their ability to detect tapeworm antigens and initiate fibrosis, but all are capable of actuating a robust fibrosis response, and H3) populations differ in their capacity to actuate peritoneal fibrosis.

If hypothesis two were to be supported, and all populations were capable of generating fibrosis, we were also interested in testing whether there was variation in the timing of initiating or resolving that response. We predicted that the resistant population may have evolved the ability to respond rapidly, to limit the growth of the tapeworm early in infection. The resistant population may also be faster to recover from fibrosis, to mitigate immunopathological costs. Additionally, given the evolutionary history of the stickleback and *S. solidus*, a predominantly freshwater parasite, we predicted that the resistant phenotype would be derived and that the susceptible lake population would respond to immune challenges in a fashion similar to the ancestral marine population.

## METHODS

### Study System

*S. solidus* has a complex lifecycle where it is trophically transmitted from copepods to sticklebacks to birds (Orr *et al.* 1969; Barber & Scharsack 2010). When a fish consumes an infected copepod, the tapeworm penetrates the intestinal wall and enters the peritoneal cavity. The tapeworm then grows rapidly and becomes capable of infecting its definitive hosts when it crosses a threshold of ~50mg (Tierney & Crompton 1992). This size threshold also corresponds with an increase in the costs and symptoms associated with tapeworm infections, including putative behavioral manipulation by the tapeworm which likely increases the probability of fish being eaten by birds (Barber & Scharsack 2010). This tapeworm is known to successfully infect a variety of bird and copepod species, but specializes on threespine sticklebacks as an obligate intermediate host (Barber & Scharsack 2010; Nishimura *et al.* 2011). The genomic structure of tapeworm populations suggests that there is likely not tight coevolution between hosts and tapeworms at the level of individual lakes (Shim et al. *in prep*). This is likely due to the fact that birds are moving readily between lakes, and thus mix tapeworm populations across the landscape, though the fish themselves are quite isolated between different watersheds.

### Field survey and breeding

The following field collections were conducted with approval from the British Columbia Ministry of Forests, Lands, Natural Resource Operations and Rural Development (Fish Collection Permit NA19-457335). The sample sites were all within the historical tribal region of the Kwakwaka’wakw First Nations. Collections were approved by the University of Connecticut IACUC (protocol A18-008).

Sayward Estuary (ancestral population, 50°22’46”N, 125°56’43”W) is a breeding location for the anadramous marine stickleback. We used this population as a proxy for the phenotype of ancestral stickleback that likely colonized freshwater lakes on Vancouver Island. Sayward fish spend the majority of their life at sea and have very little natural exposure to *S. solidus*, and are highly susceptible in laboratory experiments (Weber *et al.* 2017a). Gosling Lake (susceptible population, 50°03’47”N, 125°30’07”W), on the other hand, has a consistently high infection rate of *S. solidus* with 50% to 80% of fish infected depending on the year, and infected fish often have large tapeworms (Weber et al. 2017). Another freshwater population in Roselle Lake (resistant population, 50°31’13”N, 126°59’12”), has been noted as having lower infection rates, smaller tapeworms, and higher incidence of fibrosis (*Shim unpublished data).* Infection level for this population was found to be ~40%, though this was estimated with only 44 fish captured in 2016 (Weber *et al. in prep*). Roselle and Gosling lakes are in separate watersheds and contain isolated stickleback populations with no access to the ocean and limited capacity for gene flow with other populations in their watersheds due to inhospitable outlet and inlet streams.

In the spring of 2018, we sampled 31 uninfected and 31 infected fish from Gosling Lake and 30 uninfected and 32 infected fish from Roselle Lake using unbaited minnow traps in order to quantify average tapeworm size and the frequency and severity of fibrosis. Fish were sampled as part of a larger gene expression study where we sampled the first 30 uninfected fish and then continued sampling until we had found 30 infected fish for each population. Fish were categorized as uninfected if we did not find a living tapeworm within their peritoneal cavity. We scored fibrosis in the peritoneal cavity visually as: 0 (no fibrosis), 1 (some fibrosis, organs do not move freely), 2 (fibrosis adhering organs together), 3 (organs adhered together and to the peritoneal wall), 4 (severe fibrosis, difficult to open peritoneal cavity) (see video: https://youtu.be/yKvcRVCSpWI). We weighed tapeworms on a digital scale to the nearest 0.01g; tapeworms weighing less than 0.01g were recorded as <0.01g, as this was the lower limit of our field scale, and recorded as 0.009g for summary statistics. If fish were infected with multiple tapeworms, we weighed all tapeworms within a fish together to get average parasite mass. We compared infection intensity between lakes using a general linear model (glm) with a Poisson distribution, and average tapeworm mass (converted to mg and log transformed) using a linear model. We compared the fibrosis scores of uninfected and infected fish between lakes using a cumulative link model for ordinal data, from the package “Ordinal” (Christensen 2019). To get a more accurate estimate of infection rate for the Roselle population, we also euthanized and preserved 169 randomly selected fish in ethanol, which were later dissected and scored as infected or uninfected (ethanol preservation is not conducive to scoring fibrosis).

In June 2018, we also collected fish from our three target populations for breeding. Using standard in-vitro fertilization methods for stickleback, we created families from each population in the field and transported fertilized eggs back to the lab for rearing. Fish were housed by family in two rooms at the animal care facility of the University of Connecticut. Families were often, though not always, split across multiple tanks located in both rooms. All fish were approximately 11 months old when they were injected with different immune challenges in May 2019.

### Laboratory Injection Experiment

We injected four different inoculants directly into the peritoneal cavity of fish. These included 1) 20ul of 1X phosphate buffered saline (PBS, control treatment), 2) 10ul of homogenized tapeworm protein solution + 10ul PBS (tapeworm treatment), 3) 10ul of Alum (2% Alumax Phosphate, OZ Bioscience) + 10ul PBS (alum treatment), and 4) 10ul tapeworm protein + 10ul Alum (tapeworm + alum treatment). Alum is an immune adjuvant, or mild irritant, that causes the recruitment of leukocytes that then initiate an immune response (Kool *et al.* 2012). It is a common component of vaccines, and pilot studies demonstrated that alum injections can induce a fibrosis response in the peritoneal cavity of stickleback (*Natalie Steinel, per. comm).* The tapeworm protein solution was used to test if fish could recognize and respond to tapeworm antigens. The tapeworms we used to create the tapeworm treatment came from sticklebacks collected from Farwell Lake (50°11’60”N, 125°35’27”W) on Vancouver Island in the summer of 2008 that were flash frozen and stored at −80°C. These fish were thawed on ice and dissected to recover tapeworms. We purposely chose tapeworms from a different lake, watershed, and year in order to minimize any localized genetic structure of the parasite that may influence population level responses in our experiment. Individual tapeworms were dipped in deionized water and placed in chilled 0.9× PBS. Each tapeworm was sonified on ice twice for 1 min (Branson Sonifier 150 Ultrasonic Cell Disruptor, set to level 5). Between sonification rounds, tapeworm samples were chilled on ice for 5 min. After sonification, the homogenized tapeworm solutions were spun in a chilled centrifuge (4°C) at 4000rpm for 20 min. The supernatant from each was collected and pooled. The protein concentration of this pooled tapeworm homogenate solution was measured using a Red 660 kit (G-Biosciences), and then diluted to 1mg/ml using 0.9× pbs, after which we aliquoted and stored it at −20C.

Before injection, fish were lightly anesthetized using a neutral-buffered MS-222 (50-75 mg/L). We used ultra-fine insulin syringes (BD 31G 8mm) to inject 20ul of one of our four treatments into the lower left side of the peritoneal cavity, slightly above where the end of ventral spine rests. Injections were done as shallow as possible and at an angle parallel to the fishes body to avoid puncturing any organs, while at the same time watching for visual distention of the peritoneal cavity to ensure solutions were being injected correctly. All solutions were prepared and syringes were loaded in a sterile culture hood before injection. At the time of injection, each fish was also given a small colored elastomer mark (Northwest Marine Technologies) corresponding to their treatment group inserted above and behind the left eye. During the injection procedure, fish were placed on a wet sponge and had their head and gills covered with a wet paper towel. In total, the procedure lasted less than one minute and fish were then immediately placed in an aerated recovery tank before being returned to their home tank with negligible mortality. The protocol was IACUC-approved (protocol A18-008).

We euthanized fish to measure fibrosis responses post injection at four different time points: 1, 10, 42, and 90 days. We used the 0-4 visual fibrosis scale described above, and all fish were scored by two people (AKH & LF) who were blind to treatment and population. We also quantified fish mass and length and identified sex. To the best of our ability, we spread treatments and time points across families within each population; sample sizes are provided in table 1. We note that the sample sizes for the 90 day timepoint are smaller than the other timepoints, as this time point was added after the experiment began to take advantage of excess surviving injected fish. Given this, results for this timepoint should be interpreted with caution. Throughout the experiment, there was a mortality rate of 11% (48 out of 418 fish), which did not appear to be driven by treatment or population. Mortality typically occurred several days to weeks after the injection procedure.

**Table 1.** Sample sizes of fish that were scored for fibrosis across populations, timepoints, and treatments in our laboratory injection experiment. The number of families are indicated in parentheses.

|  |  |  |  |  |  |
| --- | --- | --- | --- | --- | --- |
| **Population** | **Time Point** | **PBS** | **Tapeworm** | **Alum** | **Tapeworm+Alum** |
| Gosling<br>*Susceptible*<br>(14) | 1 Day | 11 (9) | 10 (9) | 10 (7) | 10 (8) |
|  | 10 Days | 10 (7) | 10 (9) | 10 (8) | 10 (8) |
|  | 42 Days | 11 (8) | 10 (8) | 10 (8) | 10 (8) |
|  | 90 Days | 3 (2) | 2 (1) | 8 (3) | 4 (2) |
| Roselle<br>*Resistant*<br>(12) | 1 Day | 10 (8) | 10 (7) | 10 (8) | 10 (7) |
|  | 10 Days | 11 (8) | 10 (7) | 10 (8) | 10 (7) |
|  | 42 Days | 10 (7) | 11 (7) | 10 (7) | 10 (8) |
|  | 90 Days | 3 (2) | 8 (1) | 3 (1) | 6 (1) |
| Sayward<br>*Ancestral*<br>(9) | 1 Day | 10 (6) | 10 (6) | 11 (5) | 11 (7) |
|  | 10 Days | 10 (7) | 11 (7) | 10 (6) | 10 (5) |
|  | 42 Days | 10 (5) | 11 (5) | 11 (5) | 10 (6) |
|  | 90 Days | 2 (1) | 4 (1) | 3 (1) | 3 (1) |

### Analysis of Laboratory Injection Experiment

We built linear mixed models, using the “Linear and Nonlinear Mixed Effects Models” package (Pinheiro et al. 2019). We first built models testing whether fibrosis score depends on the main effects of, and two- and three-way interactions between, population, treatment, and time. Given that the three-way and two-way interactions were all highly significant, we ran a series of smaller models on subsets of the data in order to test hypotheses concerning specific contrasts. These models tested the following three questions: 1) Within a population, does fibrosis differ between treatments at a given time point? 2) Does the fibrosis response to a particular treatment vary between populations at given time point? and 3) Within a population, does the fibrosis response to a given treatment change through time? For each of these questions, we looked at the main effect of treatment, population, or time. If this was significant, we then used Tukey’s posthoc tests to compare between groups. All models included room as a fixed effect, though it was never significant. Family was also included as a random effect, except for some models at the 90 day timepoint where there was not enough variation in family across populations. All statistics were run using RStudio (version 1.2.1335, RStudio Team 2019).

Several alternative analytical approaches were explored but were determined to not be feasible. In particular, given that fibrosis score is ordinal, we first attempted to use cumulative link mixed models (clmms) using the package “Ordinal” (Christensen 2019). However, there was not enough variation in many of our comparison for these models to run (i.e. some treatments and time points for certain populations had low variance because all individuals had zero fibrosis, or all had strong fibrosis). Given this, we chose to use a continuous approach with the understanding that this approach requires the assumption that the numerical distances between each of our fibrosis scores are equal. However, we felt that this approximation was still the best approach available for analyzing our data, and in cases where we could get clmms to run, we found very similar results that confirm our use of a continuous approach is robust. Even with continuous models, there were still some comparisons with little variation within some groups (i.e. all responses were 0) which generated an overfitted model. In these cases, we simply report the clear pattern. In our original data exploration, we found that fish size (mass or length) and sex did not influence the fibrosis response and that models were a better fit for the data when they were removed. Given this, we excluded these factors from subsequent analyses.

## RESULTS

### Field Survey

From the 169 preserved fish collected from Roselle lake, we found 6 infected fish, giving an infection rate of 3.6%, which was lower than the rate estimated in 2016 (43%) and infection rates previously estimated for Gosling lake (50-80%, over a 5 year period) (Weber *et al.* 2017a).

Comparing the infected fish that we collected from each population in the spring of 2018 (31 from Gosling and 32 from Roselle), Gosling had a significantly higher infection intensity (GOS: mean=3.87 worms, sd=5.75; RSL: mean=2.31 worms, sd=1.82, GLM: b(Roselle)=−0.51, se=0.15, p<0.001) and higher average worm mass (GOS: mean=52.61mg, sd= 84.47; RSL: mean= 32.47mg, sd= 55.03, b(Roselle)=−0.35, se=0.35, p=0.32) though this was not statistically significant, perhaps because of the limits of our field scale (many Roselle infections fell below the 0.01g threshold and were smaller by eye compared to Gosling).

In infected fish, the degree of fibrosis was significantly higher in Roselle fish compared to Gosling fish (GOS: mean=0.06, sd= 0.25; RSL: mean= 1.72, sd=1.08; CLM: b(Roselle)=4.04, se=0.85, p<0.0001; Figure 1A). It is also of note that in seven of the fish sampled from Roselle, we found small dead tapeworms that were encased in fibrosis and partially dissolved, which we have never observed in Gosling fish. When we compared uninfected fish across lakes, fibrosis scores were also significantly higher for Roselle, though the CLM would not run because there was no variation in the Gosling population (all responses were zero), thus, we compared the populations using a t-test (GOS: mean=0, sd= 0; RSL: mean= 1.17, sd=1.46; t=−4.36, df= 29, p=0.0001; Figure 1B). As we show below, the presence of fibrosis in “uninfected” Roselle fish may be a legacy of cleared previous infections. This supports other work that has demonstrated that across lakes, populations of stickleback with more fibrosis tend to have lower infection rates and smaller tapeworms (Weber *et al. in prep*).

**Figure 1.**
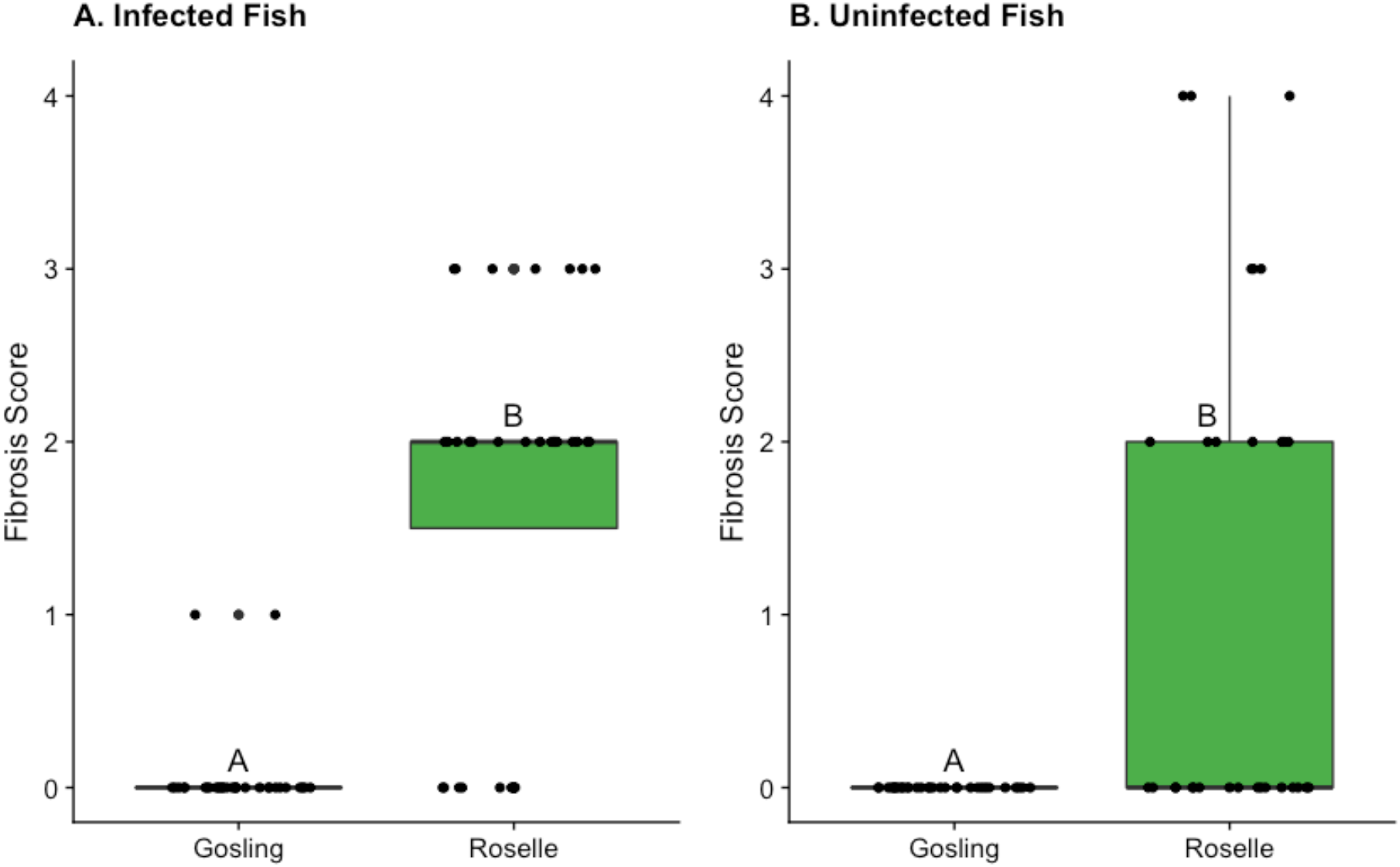
Fibrosis scores for *S. Solidus* infected (panel A) and currently uninfected (panel B) wild-caught fish from Gosling and Roselle Lakes from 2018 field data. Raw data is represented as black jittered points on the boxplot. Fish from Roselle had significantly more fibrosis in both the infected and uninfected groups relative to fish from Gosling.

### Laboratory Injection Experiment

When examining the degree of fibrosis response to injection, we found a significant three-way interaction between treatment, timepoint, and population (F_6, 362_=5.24, p< 0.0001). Once we broke this apart, all pairwise interactions were also significant (treatment*timepoint: F_3, 368_= 6.93, p<0.0001; treatment*population: F_6, 368_=2.81, p=0.01; timepoint*population: F_2, 368_=18.02, p<0.0001). To interpret these results, we used contrasts among subsets of the data to address the three questions outlined in the methods.

#### 1) Within a population, does fibrosis differ between treatments at a given time point?

For both Sayward and Gosling, fibrosis did not differ among treatments 24 hours post injections, but did differ at the 10, 42, and 90 day timepoints. For both of these populations, there was negligible fibrosis in the control and tapeworm treatment throughout and a strong fibrosis response to the alum and tapeworm+alum treatments after the first time point. Roselle produced a different pattern, where fibrosis was significantly different between treatment groups at all four time points. Roselle was the only population to show fibrosis at the first time point (in all except the control treatment) and to develop fibrosis to the tapeworm treatment, which was lower than the response to the alum and tapeworm+alum treatments. The model results are presented in Table 2, results of pairwise comparisons between treatments are summarized in Figure 2, and the least-squared means and confidence intervals for these comparisons are reported in Table S1.

**Table 2.** Statistical results (ANOVA) from models comparing fibrosis scores between treatments for a given timepoint for each population. For Sayward, 1 Day, responses were all zeros so statistical models would not run. Results of pairwise comparisons between treatments are displayed in Figure 2.

|  |  |  |  |  |
| --- | --- | --- | --- | --- |
| **Population** | **Timepoint** | **Degrees of Freedom** | **F value** | **P value** |
| Sayward<br>(*ancestral*) | 1 Day | *no response* | *no response* | *no response* |
|  | 10 Days | 29 | 78.86 | <0.0001 |
|  | 42 Days | 31 | 401.21 | <0.0001 |
|  | 90 Days | 8 | 177.00 | <0.0001 |
| Gosling<br>(*susceptible*) | 1 Day | 27 | 1.32 | 0.29 |
|  | 10 Days | 25 | 75.45 | <0.0001 |
|  | 42 Days | 23 | 33.40 | <0.0001 |
|  | 90 Days | 11 | 8.63 | 0.003 |
| Roselle<br>(*resistant*) | 1 Day | 26 | 6.00 | 0.003 |
|  | 10 Days | 29 | 41.40 | <0.0001 |
|  | 42 Days | 26 | 30.36 | <0.0001 |
|  | 90 Days | 15 | 7.86 | 0.002 |

**Figure 2.**
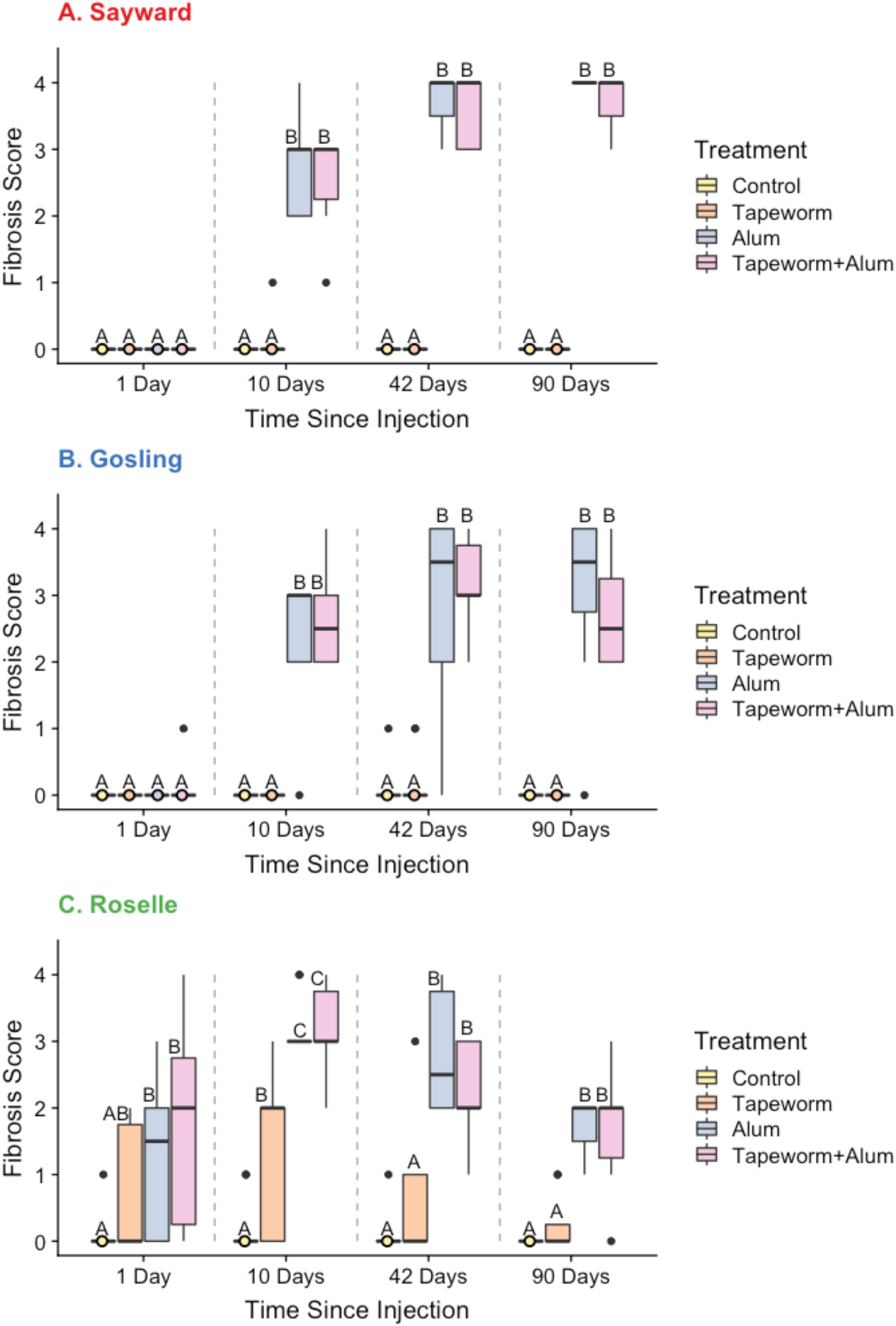
Fibrosis scores from the laboratory injection experiment for each treatment (Control, Tapeworm, Alum, & Tapeworm+Alum) at each timepoint (1, 10, 42, & 90 days point injection) for Sayward (panel A), Gosling (panel B), and Roselle (panel C) populations. Letters denote significant differences between treatments within a time point (comparisons within gray dotted lines) using Tukey’s posthoc tests (Ps < 0.05). All populations generated a strong fibrosis to the Alum and Tapeworm+Alum treatments, particularly by day 10. However, Roselle was the only population to generate a fibrosis response within 24 hours of injection and was also the only population to generate fibrosis in response to the tapeworm treatment.

#### 2) Does the fibrosis response to a treatment vary between populations at given time point?

For the control treatment, there was little fibrosis and populations did not differ at any timepoint. For the tapeworm treatment, populations differed in their fibrosis response at both 1 and 10 days, but not at 42 and 90 days, with Roselle being the only population to produce fibrosis to this treatment (through pairwise comparisons for the 1 day timepoint were not significant). For the alum treatment, fibrosis was significantly different between populations for the first timepoint but not for the rest. Specifically, Roselle was the only population to produce fibrosis within 24 hours, but Sayward and Gosling caught up to produce similar levels of fibrosis for the remaining timepoints. At 90 days, there was a trend of population differences for the alum treatment, with a decreasing response for Roselle, though small sample sizes limited our statistical power. For the tapeworm+alum treatment, fibrosis was significantly different between populations for 1, 42, and 90 day timepoints, with Roselle being the only population to respond on day 1 with a decreased response at both the 42 and 90 days relative to the other two populations. The model results are presented in Table 3, and results of pairwise comparisons are summarized in Figure 3.

**Table 3.** Statistical results (ANOVA) from models comparing fibrosis between populations for a given timepoint and treatment. For the control treatment, 1 day and 90 days, responses were all zeros so statistical models would not run. Pairwise comparisons between populations are presented in Figure 3

| Treatment | Timepoint | Degrees of Freedom | F value | P value |
| --- | --- | --- | --- | --- |
| Control (PBS) | 1 Day | *no response* | *no response* | *no response* |
|  | 10 Days | 18 | 1.71 | 0.29 |
|  | 42 Days | 17 | 0.03 | 0.97 |
|  | 90 Days | *no response* | *no response* | *no response* |
| Tapeworm Protein | 1 Day | 19 | 3.63 | 0.046 |
|  | 10 Days | 20 | 11.60 | 0.0005 |
|  | 42 Days | 17 | 1.75 | 0.20 |
|  | 90 Days | 11 | 0.79 | 0.48 |
| Alum | 1 Day | 17 | 10.37 | 0.001 |
|  | 10 Days | 19 | 3.37 | 0.06 |
|  | 42 Days | 17 | 1.95 | 0.17 |
|  | 90 Days | 14 | 3.81 | 0.34 |
| Tapeworm + Alum | 1 Day | 19 | 10.71 | 0.001 |
|  | 10 Days | 17 | 1.34 | 0.29 |
|  | 42 Days | 17 | 11.07 | 0.0008 |
|  | 90 Days | 9 | 5.72 | 0.02 |

**Figure 3.**
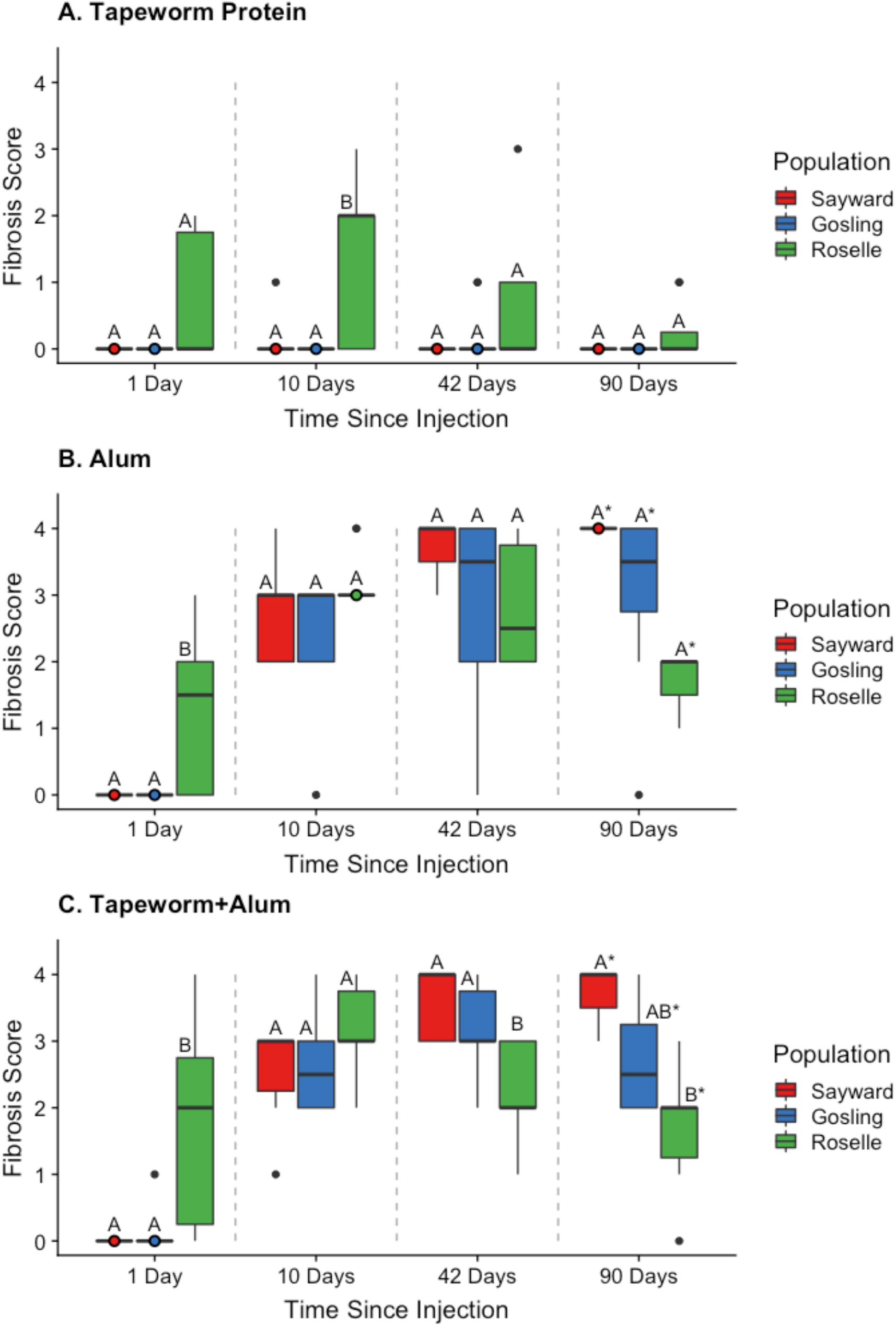
Fibrosis scores from the laboratory injection experiment for each population (Sayward, Gosling, & Roselle) at each timepoint (1, 10, 42, & 90 days post injection) for the tapeworm treatment (A panel), alum treatment (B panel), and tapeworm+alum treatment (C panel). The control treatment (PBS) is not pictured as there was little fibrosis in any of the populations. Letters denote significant differences between populations within a time point (comparisons within gray dotted lines) using Tukey’s posthoc tests (ps < 0.05). An * denotes reduced statistical power due to small sample sizes at the 90 day timepoint. The responses from Gosling and Sayward were indistinguishable, while Roselle followed a different pattern in all three treatments. Note that this is the same data as in Fig. 2, replotted to focus on population differences to a given treatment.

#### 3) Within a population, does the fibrosis response to a given treatment change through time?

For both Sayward and Gosling, fibrosis to the control and tapeworm treatments did not differ through time (negligible fibrosis at all timepoints), however, fibrosis did change through time for both the alum and the tapeworm+alum treatment, where there was no response on day 1, an increase in fibrosis from 10 to 42 days and a continuation of high fibrosis at 90 days, showing no evidence of attenuation. For the Roselle population, there was again no significant effect of time for the control and tapeworm treatments, though for the tapeworm treatment, there was a trend where fibrosis appeared to peak at ten days and then decrease at the 42 and 90 day time points. Fibrosis did significantly change through time for the other treatments, where fibrosis was detected at the first time point, peaked at ten days, and by 90 days had decreased, suggesting attenuation of the response. The model results are presented in Table 4, and results of pairwise comparisons between populations are summarized in Figure 4.

**Table 4.**
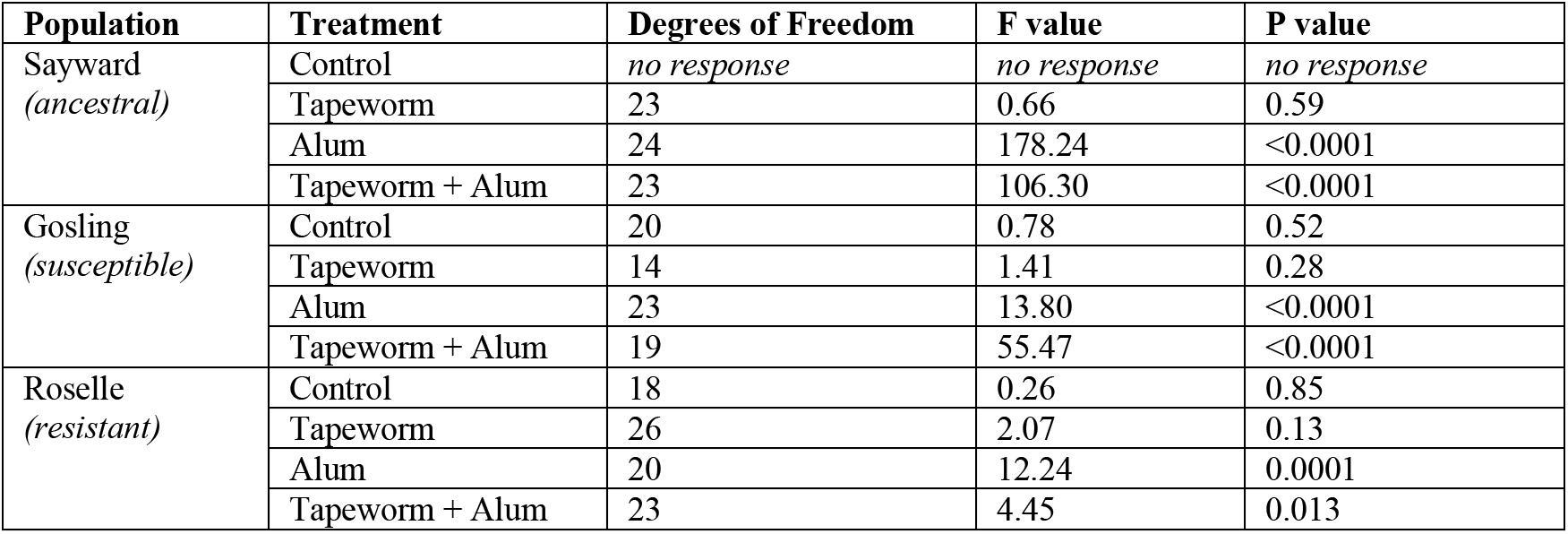
Statistical results (ANOVA) from models comparing fibrosis scores for a given treatment through time for each population. For Sayward, 1 day, responses were all zeros so statistical models would not run. Pairwise comparisons between timepoints are displayed in Figure 4.

**Figure 4.**
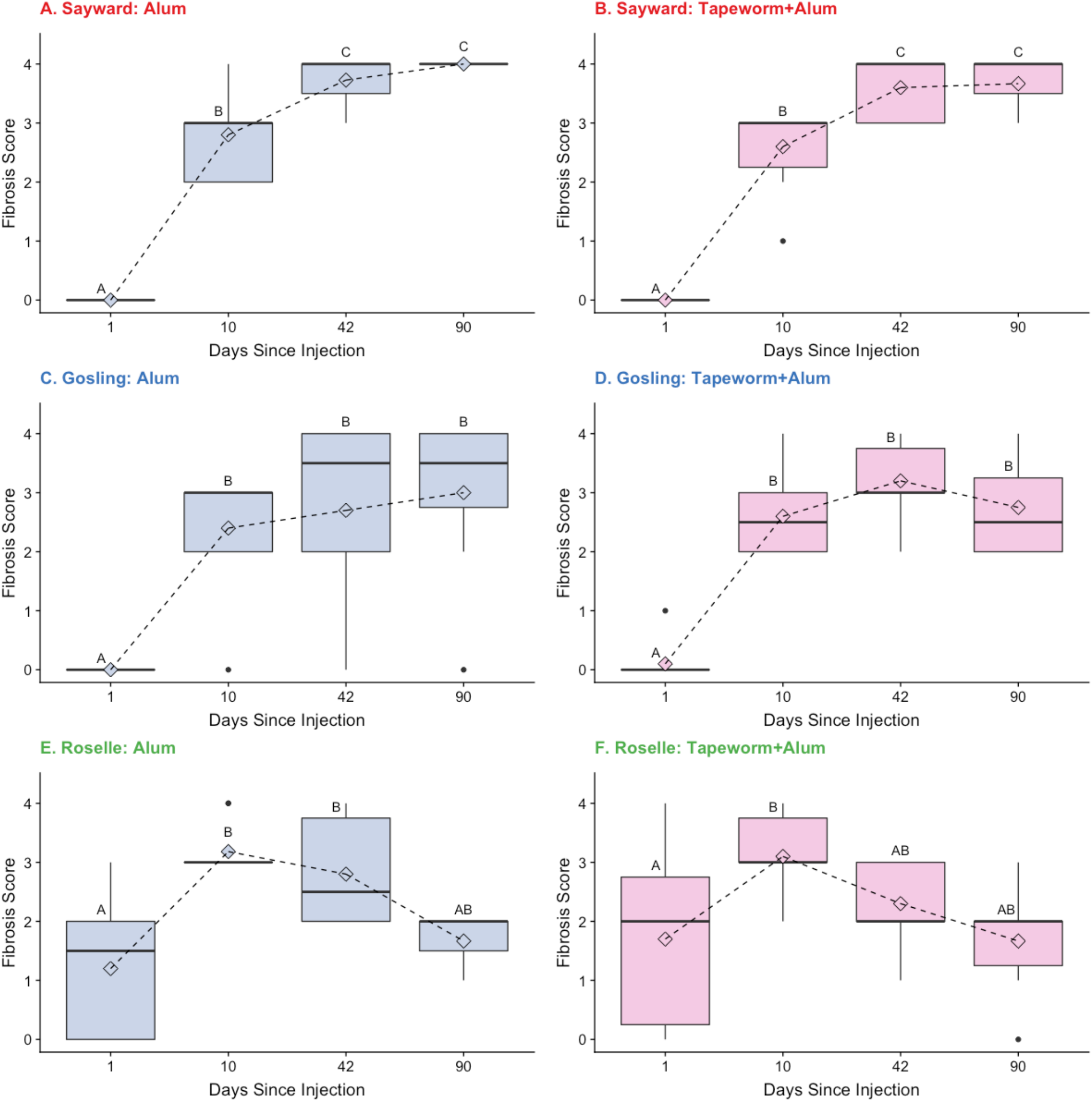
Fibrosis scores from Alum and Tapeworm+Alum injection treatments (PBS and tapeworm treatments not pictured) through time (1, 10, 41, and 90 days post injection) for Sayward (panels A & B), Gosling (panels C & D) and Roselle (panels E & F). Letters denote significant differences between timepoints using Tukey’s posthoc tests (Ps < 0.05). Roselle initiated fibrosis earlier, within 24 hours, and then showed evidence of a decreased response (attenuation) in the last time point(s) of the experiment, unlike Sayward and Gosling which either increased or maintained a high level of fibrosis after day 10.

## DISCUSSION

Striking differences in parasite resistance between related groups, even to the same parasite, is common in nature, and yet, we often lack an understanding of the immunological mechanisms that generate this important variation. While many evolutionary studies emphasize the broad outcomes of infection (e.g., success or failure or parasite load), we instead take a stepwise approach that breaks up different components of the host response through time to better understand resistance evolution in populations of threespine stickleback. In this study, we compare an ancestral marine population and two lake populations that differ in their resistance response to a freshwater tapeworm. Using a novel vaccination assay, we set out to test if H1) variation in resistance was being driven only by the environment, H2) variation was due to population-level differences in the ability to detect and respond to tapeworm antigens, or H3) variation was due to population-level differences in the ability to generate a strong fibrosis response in the peritoneal cavity. We also tested for variation in the rate of initiation and resolution of fibrosis, with the prediction that selection may have favored rapid initiation and quicker resolution in the resistant population, to maximize efficacy of the early response and mitigate long-term costs of fibrotic pathology.

Our results demonstrate that while all populations were capable of producing a robust fibrosis response in their peritoneal cavity, only the resistant population was able to respond to the tapeworm antigen alone, supporting H2. This suggests that population-level variation in tapeworm resistance is likely driven by variation in parasite recognition that then leads to the initiation of fibrosis. But, all populations retain the capacity to actuate a fibrotic response, which we subsequently showed is a highly conserved immune response across teleost fish (Vrtílek & Bolnick *in prep*). We also found that the resistant population produced fibrosis faster, within 24 hours of injection, and then attenuated that response during our experimental period, which was not the case for the other populations. Finally, by comparing these populations, our findings suggest that the more resistant phenotype (Roselle), which can recognize tapeworm antigens, respond rapidly (within 24 hours) and then attenuate that response, is the derived state. This conclusion is supported by the observation that the susceptible population (Gosling) largely matches the responses of the ancestral outgroup represented by marine fish (Sayward).

Our findings, which suggest that parasite detection mechanisms are important in generating variation in parasite resistance across populations, fits well into recent work that has highlighted the importance of identity signatures in host-parasite coevolution (Spottiswoode & Busch 2019). These self/non-self-recognition systems are known to evolve rapidly and can lead to varying outcomes in the arms race between hosts and parasites in different contexts. Given this, they may be a widespread mechanism generating rapid variation in infection outcomes between individuals and populations, even under short evolutionary timescales (Radwan *et al.* 2020). For large macroparasites, such as helminths, it is known that dendritic cells play a central role in recognition and initiation of an immune response (Motran *et al.* 2018). However, the transition between successful parasite recognition and the initiation of an immune response with helminth infections is particularly tricky to study, as helminths are well known to secrete a suite of immunomodulatory molecules that suppress and shift host immune responses (Coakley *et al.* 2016; Maizels *et al.* 2018; Motran *et al.* 2018). By using homogenized tapeworm tissue, our experimental design mitigates active interference by the tapeworm, allowing us to test if hosts can recognize and respond to tapeworm antigens without the accompanying immunomodulation of a live infection.

The mechanism of this between-population divergence in recognition is unclear at present. A commonly invoked candidate gene for helminth recognition, Major Histocompatibility Complex IIb, exhibits widespread covariation between allelic composition and macroparasite infection in stickleback and other vertebrates (Bernatchez & Landry 2003; Kurtz *et al.* 2004; Eizaguirre *et al.* 2010). But, a recent survey of six stickleback populations found no associations between stickleback MHCIIß genotype and *S. solidus* prevalence, though many other macroparasites’ intensity was correlated with MHCIIß alleles (Stutz & Bolnick 2017). A larger unpublished study of 25 populations also found little evidence for MHCIIß associated with *S. solidus* (Peng et al. *in prep*). Moreover, QTL mapping of Gosling Lake versus another high-fibrosis low-infection population (Roberts Lake) found QTLs for fibrosis, infection, and cestode growth but none of these contained MHC loci (Weber et al. *in prep*). Then there is the consideration that MHCIIß immune responses typically involve the adaptive immune system which takes time to initiate a response (especially in naïve fish that lack any prior immune memory, and in ectotherms in cold water) (Wegner *et al.* 2007). In contrast, we see the start of a fibrotic response to cestode protein within 24 hours. It is more likely that the populations differ in receptors involved in detecting *S.solidus* antigens, which then initiate innate responses, or innate-like lymphoid cell (ILC) responses. Thus, our findings concerning the role of recognition, and the speed of the response, help narrow the ongoing search of possible immunological mechanisms and genes. Our findings indicate that such genes or mechanisms likely vary across populations.

Our results also suggest that within the resistant population, there has been selection on the timing of both initiating and resolving the fibrosis response. Biologically, this makes sense because if fish can respond early, while tapeworms are small, they are likely to be more successful in clearing the infection. This could explain what we witnessed in the field, where small dead tapeworms were trapped in fibrosis. Even if fibrosis does not successfully kill tapeworms, early initiation could still limit their growth (Weber *et al. in prep*), allowing fish to avoid many of the negative consequences imposed by large tapeworms. If fish can successfully clear infection, resolving the fibrosis response is likely adaptive, as we hypothesize that this response is likely costly to maintain and may impact growth, swimming ability, and reproduction. It is clear from work in other systems that variation in the timing, and not necessarily differences in the magnitude, of the immune response can lead to striking differences in infection outcomes (Duneau *et al.* 2017). It will be informative for future work to extend the timeline of this experiment, to see what is occurring after 90 days and if a strong fibrosis response can fully resolve back to a normal state. We were also limited in our ability to detect some of these patterns by our small sample sizes, particularly at the last timepoint. In the marine and susceptible populations, which were unable to recognize tapeworm protein, the machinery to produce a strong fibrosis response was clearly still there, but there has likely not been strong selection on the timing of initiation or resolution of that response, as it fails at an early step in the sequential chain of events-parasite recognition. In all three populations, however, significant fibrosis (greater than the control treatment) persisted to 90 days after a single immune challenge. This highlights the potential long-term cost of mounting such a response, particularly in a short-lived fish. It also explains why we observe fibrotic individuals without *S.solidus* in the wild, while there is negligible spontaneous fibrosis of immunologically naïve fish. Stickleback apparently can mount a successful fibrotic response that eliminates the tapeworm, but which then persists after the infection is cleared.

Our study worked to isolate the genetic contribution to variation in the response to infection across our study populations, by raising all fish in the same laboratory conditions and injecting them with different immune challenges (removing, for example, variation in exposure or resources). This approach can be informative, as it allows for the isolation of genes from ecology, but translating our results back to wild populations must be done with some caution. It is clear that ecology can interact with genetics to play an important role in shaping host responses to infection and generating variation between populations (Hawley & Altizer 2011; Leung *et al.* 2018). Additionally, using the same tapeworm extract allows us to easily compare across populations, but does not address how a live tapeworm might interfere with the fibrosis response (Steinel & Bolnick 2018; Piecyk *et al.* 2019, Fuess et al. *in prep*), and beyond that, how variation in the tapeworms themselves might further shape host responses (e.g. gene-for-gene epistasis between species). We also acknowledge that our study is limited to only three populations, and it will be informative for future work to determine if the same patterns exist between other susceptible and resistant lake populations. Ongoing gene expression work using samples collected from this experiment will also provide more insights into the genetic mechanisms underlying these patterns and will give more detail about what the immune response is doing beyond the visual fibrosis score used here.

By taking a stepwise approach to isolate different stages of the host response to infection, we were able to uncover clear evidence that differences arising at key stages, namely parasite recognition and the timing of fibrosis initiation, are likely driving variation in parasite resistance between closely related populations of threespine stickleback. Differences in fibrosis clearance are also likely to play a role in adaptive tuning of the costs of initiating the response. Applying this approach, of partitioning variance across infection stages, in a variety of wild systems is likely to provide a more nuanced understanding of the mechanisms generating variation in parasite resistance and insight into host-parasite coevolution.

## ACKNOWLEDGMENTS

Thanks to J. Salguero for help with fieldwork, and to J. Weber and N. Steinel for help with fieldwork, crossing fish, and for their advice and support during the laboratory experiment. We also thank S. De Lisle for his help with the experimental design and early planning of the laboratory experiment. Thanks to members of the Bolnick and Snell-Rood labs for providing helpful feedback on this manuscript. Funding and support for this work was provided by a James S. McDonnell Foundation Postdoctoral Fellowship to AKH, an American Association of Immunologists Intersect Postdoctoral Fellowship to LEF, University of Connecticut (startup to DIB), and NIH NIAID grant to 1R01AI123659-01A1 to DIB.

## AUTHOR CONTRIBUTIONS

AKH, KCS, and DIB carried out all field work and KCS dissected preserved fish from Roselle, AKH, LEF, and DIB designed the laboratory experiment. AKH, LEF, MLK, MFM, JMM carried out the laboratory experiment. AKH performed all analyses, made all figures, and wrote the manuscript. All authors provided feedback on the manuscript.

## DATA ACCESSIBILITY

Upon acceptance, all data for this manuscript will be made publicly available via Dryad and the data DOI will be included in the article. All code for analyses and figures will also be made publicly available.

## Supplement

**Table S1.** Least squared means and confidence intervals for fibrosis response results from the laboratory injection experiment. Broken up by time, population, and treatment. Exact values of least squared means and confidence intervals varied slightly depending on how models were constructed (question 1, 2, and 3 from the main text), but results were consistent.

|  | 1 Day |  | 10 Days |  | 42 Days |  | 90 Days |  |
| --- | --- | --- | --- | --- | --- | --- | --- | --- |
|  | LSM | CI | LSM | CI | LSM | CI | LSM | CI |
| **Sayward (Ancestral Population)** |  |  |  |  |  |  |  |  |
| PBS | 0 | *na* | -0.02 | -0.43, 0.40 | 0.02 | -0.25, 0.28 | 0.00 | *na* |
| Alum | 0 | *na* | 2.80 | 2.39, 3.21 | 3.71 | 3.46, 3.97 | 4.00 | *na* |
| Tapeworm | 0 | *na* | 0.09 | -0.30, 0.48 | -0.02 | -0.27, 0.24 | 0.00 | *na* |
| Tapeworm+Alum | 0 | *na* | 2.62 | 2.20, 3.03 | 3.62 | 3.35, 3.88 | 3.67 | *na* |
| **Gosling (Susceptible Population)** |  |  |  |  |  |  |  |  |
| PBS | -0.01 | -0.12, 0.10 | -0.10 | -0.54, 0.33 | 0.10 | -0.50, 0.70 | -0.31 | -3.44, 7.83 |
| Alum | 0.00 | -0.11, 0.11 | 2.30 | 1.87, 2.73 | 2.70 | 2.07, 3.32 | 2.98 | -1.80, 7.75 |
| Tapeworm | -0.00 | -0.11, 0.11 | -0.03 | -0.45, 0.39 | 0.17 | -0.47, 0.80 | 0.35 | -9.55, 10.25 |
| Tapeworm+Alum | 0.11 | -0.00, 0.22 | 2.65 | 2.22, 3.08 | 3.23 | 2.59, 3.86 | 2.85 | -3.94, 9.64 |
| **Roselle (Resistant Population)** |  |  |  |  |  |  |  |  |
| PBS | 0.04 | -0.72, 0.81 | 0.15 | -0.41, 0.71 | 0.15 | -0.38, 0.69 | 0.00 | -5.02, 5.02 |
| Alum | 1.24 | 0.47, 2.00 | 3.18 | 2.63, 3.74 | 2.73 | 2.19, 3.27 | 1.67 | -3.36, 6.69 |
| Tapeworm | 0.69 | -0.09, 1.47 | 1.39 | 0.81, 1.98 | 0.59 | 0.07, 1.11 | 0.25 | -2.83, 3.33 |
| Tapeworm+Alum | 1.69 | 0.92, 2.47 | 3.13 | 2.55, 3.72 | 2.320 | 1.76, 2.88 | 1.67 | -1.88, 5.22 |

